# Dynamics of pax7 expression during development, muscle regeneration, and *in vitro* differentiation of satellite cells in the trout

**DOI:** 10.1101/2023.07.19.549701

**Authors:** Cécile Rallière, Sabrina Jagot, Nathalie Sabin, Jean-Charles Gabillard

## Abstract

Essential for muscle fiber formation and hypertrophy, muscle stem cells, also called satellite cells, reside beneath the basal lamina of the muscle fiber. Satellite cells have been commonly localized by the expression of the Paired box 7 (Pax7) due to its specificity and the availability of antibodies in tetrapods. In fish, the identification of satellite cells remains difficult due to the lack of specific antibodies in most species. Based on the development of a highly sensitive *in situ* hybridization (RNAScope^®^) for *pax7*, we showed that *pax7*^+^ cells were detected in the undifferentiated myogenic epithelium corresponding to the dermomyotome at day 14 post-fertilization. Then, from day 24, *pax7*^+^ cells gradually migrated into the deep myotome and were localized along the muscle fibers and reach their niche in satellite position of the fibres after hatching. Our results showed that 18 days after muscle injury, a large number of *pax7*^+^ cells accumulated at the wound site compared to the uninjured area. During the *in vitro* differentiation of satellite cells, the percentage of *pax7*^+^ cells decreased from 44% to 18% on day 7, and some differentiated cells still expressed *pax7*. Taken together, these results show the dynamic expression of *pax7* genes and the follow-up of these muscle stem cells during the different situations of muscle fiber formation in trout.

## Introduction

Skeletal muscle consists of multinucleated cells called myofibers that result from the fusion of muscle precursors cells called myoblasts. Myofibers formation (hyperplasia) occurs during the embryonic and fetal period and subsequently is restricted to muscle regeneration in adult mammals, whereas in large growing fish, hyperplasia persists throughout the post-larval period^1^. Myoblasts proliferate and differentiate into myocytes, which fuse to form multinucleated myotubes, and mature into functional myofibers^2^. After birth, the myoblasts are derived from the activation and proliferation of adult muscle stem cells, called satellite cells because of their position on the surface of the myofiber beneath the surrounding basal lamina^3^. In mammals and birds, satellite cells have been shown to originate from the undifferentiated myogenic dermomyotome epithelium surrounding the primary myotome and migrate through the deep myotome to reach their final position beneath the basal lamina^4,5^.

Initially, satellite cells were identified by electron microscopy, based on their anatomical location between the myofiber membrane and the basal lamina, in numerous mammalian species, but only in a few fish such as carp^6^ and zebrafish^7^. Subsequently, satellite cells have been localized by the expression of many specific genes such as desmine, M-Cadherin, myf-5, etc^2^. Among these satellite cell markers, the Paired box 7 (Pax7) is the most widely used marker due to its specificity and the availability of antibodies^8^. In mammals, *pax7* is expressed in satellite cells and myoblasts, and is essential for satellite cell survival, cell fate and self-renewal^9–11^. In zebrafish, *pax7a* and *pax7b* are expressed by satellite cells and are required for white muscle regeneration^7^. In fish, the identification of satellite cells remains difficult due to the lack of specific antibodies in most species. In zebrafish, Pax7 immunolabeling partially recapitulates the pattern of *pax7* expression visualized in a transgenic line expressing GFP under the control of the *pax7a* promoter^12^. Surprisingly, very rare Pax7 positive cells were identified in the white muscle of adult zebrafish ^7^. Using a heterologous antibody, Steinbacher et al (2007) were able to localize Pax7 positive cells in the myotome of brown trout embryos^13^. In giant danio (*Devario cf. Aequipinnatus*), Pax7 immunolabeling shows that almost all the cells (>99%) freshly extracted from white muscle are positive for Pax7, whereas Pax7-positive cells were no longer detected 24h later^14^. In rainbow trout, despite numerous attempts with appropriate controls, we were unable to obtain a specific *pax7* signal in muscle or *in vitro* using the same antibody and protocol as Froehlich et al., (2013). Furthermore, using *in situ* hybridization, *pax7* expression was readily detected in the dermomyotome during embryonic development^15,16^. Therefore, the dynamics of migration of *pax7-* positive cells to their final niche in fish remains unknown.

Based on the development of a highly sensitive *in situ* hybridization technique ^17^ (RNAScope^®^), this work aims to describe the dynamics of *pax7* expression during embryonic and larval stages, muscle regeneration and *in vitro* satellite cell differentiation in rainbow trout.

## Materials and methods

### Animals

All the experiments presented in this article were developed in accordance with the current legislation governing the ethical treatment and care of experimental animals (décret no. 2001-464, May 29, 2001), and the muscle regeneration study was approved by the Institutional Animal Care and Use Committee of the INRAE PEIMA (Pisciculture Expérimentale INRAE des Monts d’Arrée, Sizun, Britany, France; DOI : 10.15454/1.5572329612068406E12). Fish for cell culture were reared at the LPGP fish facility (DOI : 10.15454/45d2-bn67) approved by the Ministère de l’Enseignement Supérieur et de la Recherche (authorization no. D35-238-6).

### Muscle regeneration experiment

This experiment was carried out at the INRAE facility PEIMA. Briefly, 1530 ± 279 g rainbow trout (*O. mykiss*) were anesthetized with MS-222 (50 mg/l) and using a sterile 1.2 mm needle, the left side of each fish was injured by a puncture behind to the dorsal fin and above the lateral line. The right side of each fish was used as a control. White muscle samples were collected from both sides (in the injured region and contralateral) at 0, 1, 2, 4, 8, 16, and 30 days post-injury using a sterile scalpel after euthanasia by an overdose of MS-222 (200 mg/l). The obtained samples were properly stored in liquid nitrogen until further processing for gene expression analyses. No infection was detected during the experiment and the survival rate was 100%. Another muscle regeneration experiment was performed on 30 g trout and samples were collected 18 days after injury and fixed with 4% paraformaldehyde overnight at 4°C and embedded in paraffin.

### Trout satellite cell culture

Satellite cells from trout white muscle (200-250g body weight) were cultured as previously described^18^. Briefly, muscle tissue was mechanically and enzymatically digested (collagenase, C9891 and trypsin, T4799) prior to filtration (100µm and 40µm). Cells were seeded on poly-L-lysine and laminin precoated Lab-Tek™ II Chamber Slide™ (#154534, ThermoScientific, 8 wells) at a density of 80,000 cells/ml and incubated at 18°C in DMEM (D7777) with 10% fetal bovine serum to stimulate cell proliferation and differentiation. Cells were washed twice with PBS and fixed with ethanol/glycine (pH2) from day 2 to day 7 of culture.

### RNA extraction, cDNA synthesis, and quantitative PCR analyses

Total RNA was extracted from 100 mg of muscle using TRI reagent (Sigma-Aldrich, catalog no. T9424), and its concentration was determined using the NanoDrop ND-1000 spectrophotometer. One µg of total RNA was used for reverse transcription (Applied Biosystems kit, catalog no. 4368813). Trout *pax7a1* (forward, 5’-TGGGACTACGATTTATAGTTCGATTT-3’; and reverse, 5’- TTCTTACTCGCGCAAAGTCC-3’), pax7a2 (forward, 5’-TGGGACTACGATTTTATTGTCTCC-3’; and reverse, 5’-TCGTGCAAAGTCCAGACAAG-3’), and pax7b primers (forward, 5’- CGTCAAAACAATTACCACAAACA-3’; and reverse, 5’-AAAGACGACTGCATTCTACAGC-3’) were designed at exon-exon junctions to avoid amplification of genomic DNA. Secondary structure formation in the predicted PCR product were determined with the mFOLD software. Quantitative PCR analyses were performed with 5 µl of cDNA using SYBR© Green fluorophore (Applied Biosystems), according to the manufacturer’s instructions, with a final concentration of 300 nM of each primer. The PCR program used was as follows: 40 cycles of 95 °C for 3 s and 60 °C for 30 s. The relative expression of target cDNAs within the sample set was calculated from a serial dilution (1:4–1:256) (standard curve) of a cDNA pool using StepOneTM software V2.0.2 (Applied Biosystems). Real-time PCR data were then normalized to elongation factor-1 alpha (eF1α) gene expression as previously described^19^.

### In situ hybridization

For the detection of *pax7* expression in trout embryos, samples were fixed with 4% paraformaldehyde overnight at 4°C and embedded in paraffin. Cross-sections (7 µm) of muscle were then cut using a microtome (HM355; Microm Microtech, Francheville, France) and *in situ* hybridization was performed using RNAscope Multiplex Fluorescent Assay v2 (Bio-Techne #323100) according to the manufacturer’s protocol. Sections were briefly baked at 60°C for 1 hour, deparaffinized, and air dried. After 10 min in hydrogen peroxide solution (Bio-Techne #322335), sections were treated with 1X Target Retrieval (#322000; Bio-Techne) for 15 min at 100°C, followed by 25 min with Protease Plus solution (#322331; Bio-Techne) at 40°C. Due to the presence of three *pax7* genes in the trout genome^20^, we designed a set of probes targeting *pax7a1*, *pax7a2* and *pax7b* mRNA (see supplementary Fig 1). This probe set was hybridized for 2 hours at 40°C. All steps at 40°C were performed in an Bio-Techne HybEZ II hybridization system (#321720). The RNAscope Multiplex Fluorescent Assay allows simultaneous visualization of up to three RNA targets, with each probe being assigned to a different channel (C1, C2 or C3) with its own amplification steps. For embryo sections, *pax7* transcripts were targeted with the fluorescent dyes Opal 520 (#FP1487001KT; Akoya Biosciences). At the end of the *in situ* hybridization protocol, embryo sections were incubated with a primary antibody against salmon Collagen 1 (#520171; Novotec, France) and then with a secondary antibody conjugated with an Alexa Fluor 594 fluorescent dye (#A11005; ThermoFisher). Nuclei were counterstained with DAPI (0.5µg/ml) and sections were mounted with ProlongGold (P36930, Invitrogen).

Detection of *pax7* expression was also performed using the chromogenic RNAscope® 2.5HD detection reagent RED kit (#322360; Bio-Techne) in sections of trout embryos and white muscle from 4 g and 9 g fish. As previously described, samples were fixed in 4% paraformaldehyde overnight at 4°C, embedded in paraffin and cross sections (7 µm) were made. Pax7 *in situ* hybridization was performed as described above and chromogenic detection using Fast Red, was performed according to the manufacturer’s protocol. Nuclei were counterstained with DAPI (0.5 µg/ml) and sections were mounted with EcoMount (EM897L, Biocare medical). In this chromogenic RNAscope assay, the red signal can be observed with either a white light or fluorescence microscope.

For multiplex RNAscope *in situ* hybridization, fixed cells were hybridized using the RNAscope Multiplex Fluorescent Assay v2 (Bio-Techne #323100) according to the manufacturer’s protocols. (see above). *Pax7* and *myomixer* transcripts were detected using the fluorescent dyes Opal 620 (Akoya Biosciences #FP1495001KT) and Opal 520 (Akoya Biosciences #FP1487001KT), respectively. Nuclei were counterstained with DAPI (0.5 µg/ml) and cells were mounted with Mowiol^®^.

Wheat germ agglutinin (WGA) conjugated with Alexa 488 (Molecular Probes # W11261) was used to visualize connective tissue and basal lamina^21^. WGA is a lectin-based molecule specifically binds to N-acetyl-D-glucosamine and N-acetylneuraminic acid (sialic acid) residues. After four washes with PBS, sections were stained with WGA (5 µg/ml) for 3 hours at room temperature.

All the images were taken with a Canon digital camera coupled to a Canon 90i microscope.

### Automated quantification of pax7^+^ and mmx^+^ cells

To automatically quantify the number of cells expressing *pax7*, *mmx* or *pax7* and *mmx*, we adapted a macro-command^22^ on Fiji software^23^ to quantify puncta corresponding to the RNAscope labeling, per cell. Due to the presence of one or two puncta in some cells with the negative control (DapB probe #504081, Bio-Techne), a cell was considered positive when at least 3 puncta were detected. Our quantification method is available on https://gitlab.univ-nantes.fr/SJagot/fijimacro_rnascopecells.

### Statistical analyses

Data were analyzed using the non-parametric Kruskal–Wallis rank test followed by the post-hoc Dunn test. All analyses were performed with the R statistical package (version 4.2.1).

## Results

### Pax7 positive cells were detected in the deep myotome before hatching

To determine the stage of appearance of *pax7*^+^ cells in the deep myotome, we performed highly sensitive *in situ* hybridization (RNAscope technology) with *pax7* probes on cross sections of 14-28 dpf embryos (Fig 1). At day 14, *pax7*^+^ cells were detected in the neural tube and in the undifferentiated myogenic epithelium corresponding to the dermomyotome. At day 19, no *pax7*^+^ cells were detected in the fully differentiated deep myotome, excepted near to the horizontal septum, where rare *pax7*^+^ cells were observed. Two days later (D21), the majority of *pax7* expressing cells *were* still localized in the dermomyotome, and very rare *pax7*^+^ cells appeared scattered throughout the deep myotome. From day 24 to day 28, *pax7*^+^ cells were readily detected in the deep myotome, while a slight decrease in *pax7* expression in the dermomyotome epithelium was observed. *In situ* hybridization of longitudinal sections of 28-days-old embryos showed that *pax7*^+^ cells were localized along the muscle fibers of the deep myotome (Fig 2) and not in the myoseptum.

**Fig 1.**
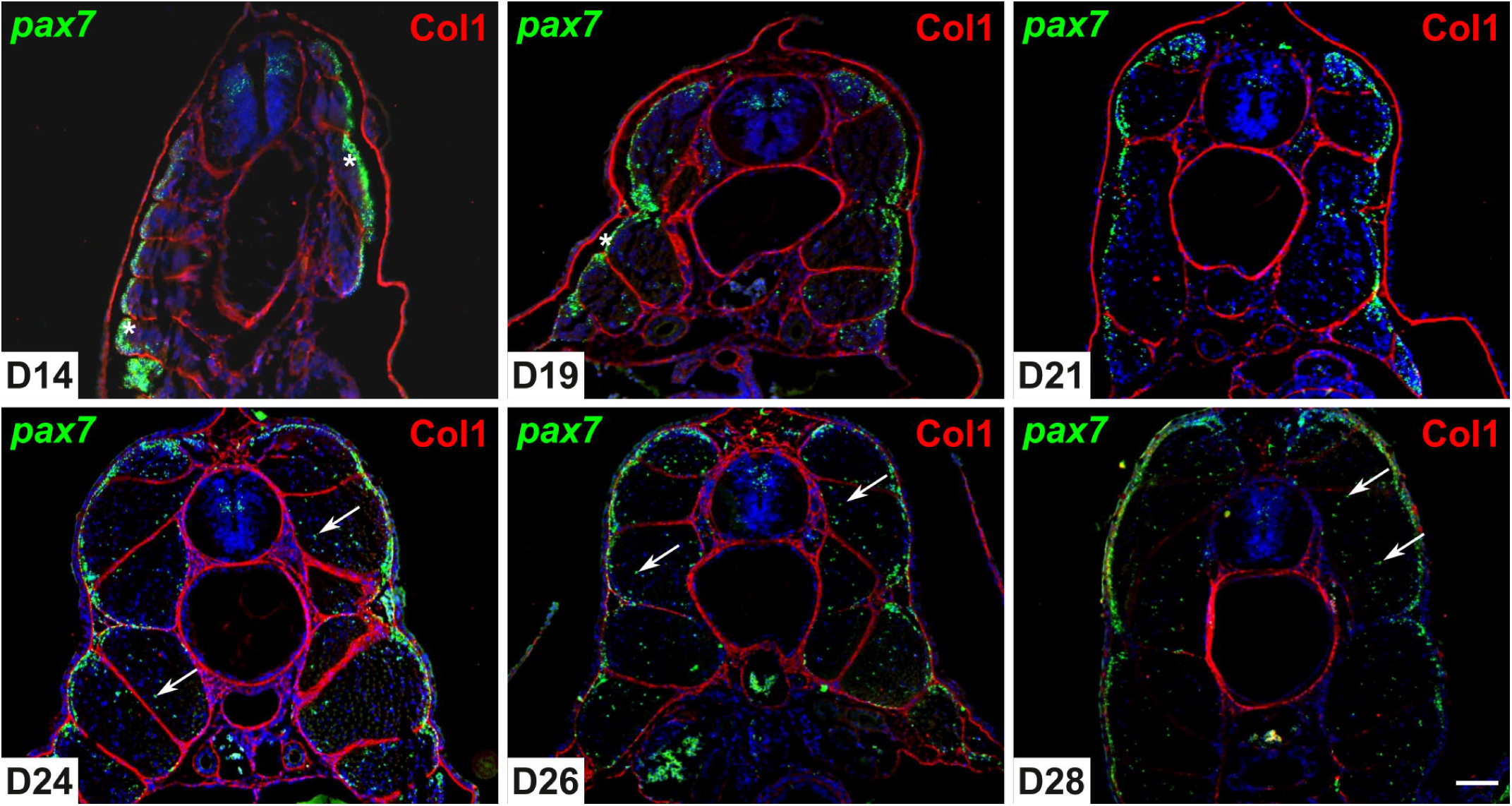
*Pax7* positive cells migrate into the deep myotome before hatching. Transverse sections (7 µm) of trout embryos were analyzed by *in situ* hybridization for *pax7* (green) and by immunofluorescence for Collagen 1 (red) at days 14, 19, 21, 24, 26, and 28 post fertilization. Asterisks indicate the dermomyotome-like epithelium and arrowheads indicate *pax7*^+^ cells in the deep myotome. The nuclei are counterstained with DAPI and the scale bar corresponds to 100 µm.

**Fig 2.**
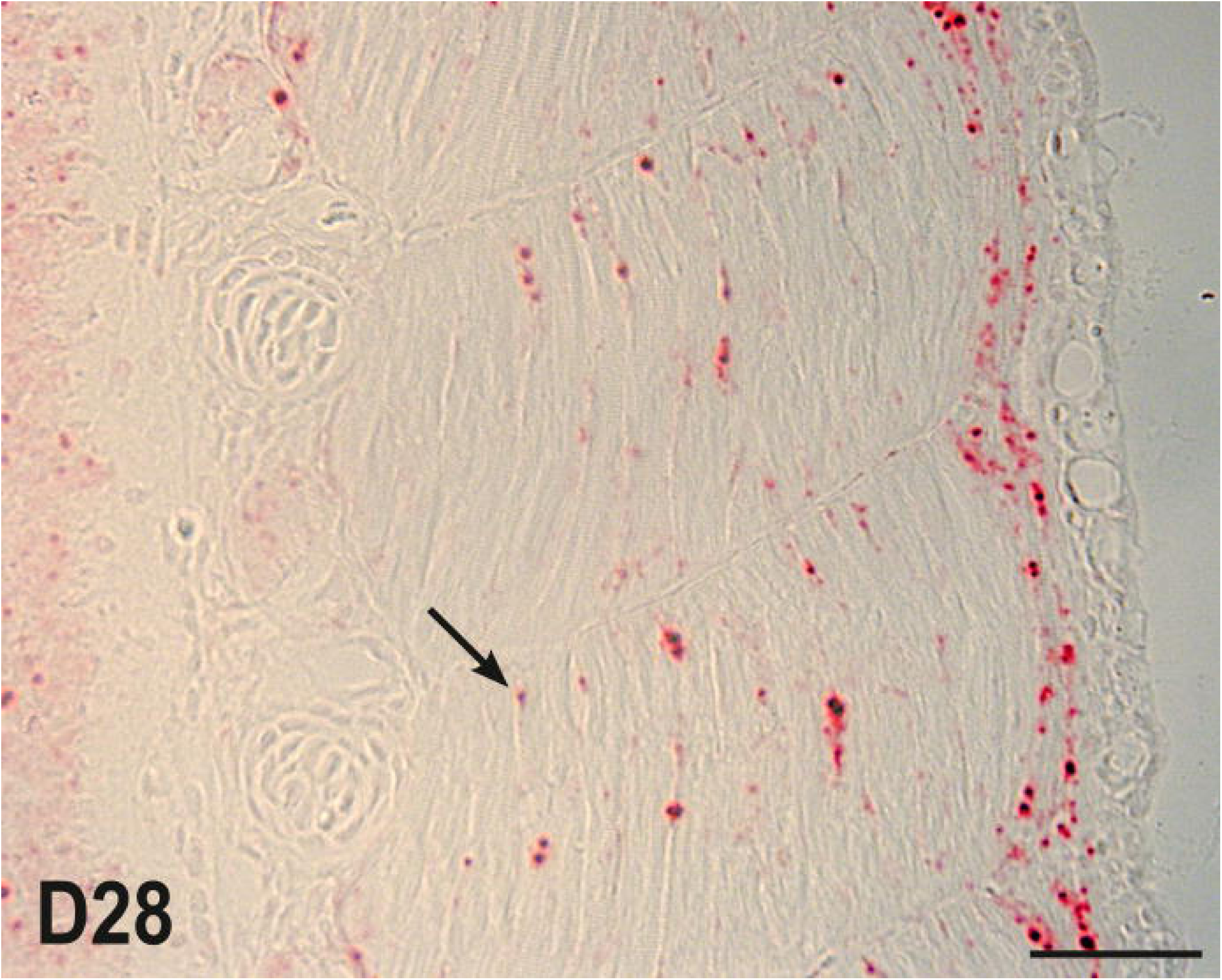
*Pax7* positive cells are positioned along muscle fibers before hatching. *Pax7* expression was analyzed by chromogenic *in situ* hybridization for *pax7* on longitudinal sections of 28-dfp trout embryos. Arrowheads indicate *pax7*^+^ cells in the deep myotome, adjacent to muscle fibers. The scale bar corresponds to 50 µm.

The final location of satellite cells is beneath the basal lamina of the muscle fibers. To determine when *pax7*^+^ cells reach their niche, we stained the basal lamina with Alexa 488-conjugated wheat germ agglutinin. In Fig 3, *pax7*^+^ cells were easily detected at day 37, when the basal lamina was very thin and did not completely surround the muscle fibers. At day 47 (100mg), the basal lamina was thicker and surrounded most of the fibers and some of the *pax7*^+^ cells. At day 112 (4g) and 136 (9g), all *pax7*^+^ cells were located beneath the basal lamina surrounding the muscle fibers.

**Fig 3.**
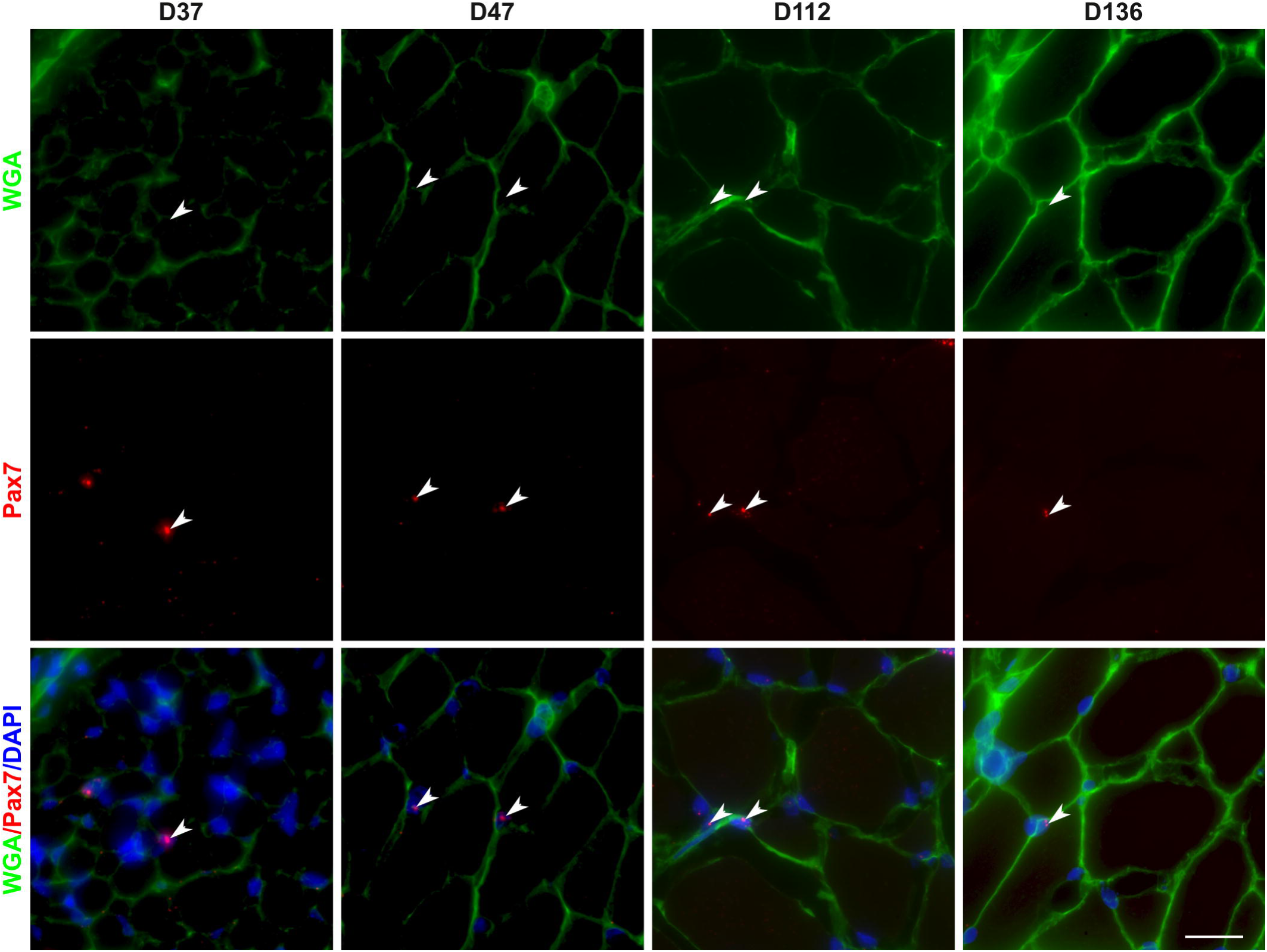
Localization of *pax7* positive cells in the muscle stem cell niche within the white muscle of trout. Transverse sections (7 µm) of trout larvae (37, 47, 112 and 136 days post fertilization) were analyzed by *in situ* hybridization for *pax7* (red) and the extracellular matrix was stained with Alexa 488-conjugated wheat germ agglutinin (green). Arrowheads indicate *pax7*^+^ cells in the deep myotome. The nuclei are counterstained with DAPI and the scale bar corresponds to 25 µm.

### Pax7 positive cells accumulate at the lesion site during regeneration

In vertebrates, muscle regeneration requires the activation, proliferation and differentiation of satellite cells (*pax7^+^)*. To determine whether the expression of *pax7* genes (*pax7a1*, *pax7a2* and *pax7b)* are upregulated during muscle regeneration in trout, we examined the kinetics of regeneration at 0, 1, 2, 4, 8, 16, and 30 days after injury. QPCR analysis showed that the expressions of *pax7a1*, *pax7a2* and *pax7b* were all upregulated after injury with a maximal expression at day 8 with an increase of 8-, 12- and 5-fold, respectively. Thereafter, the expression of *pax7a1* and *pax7a2* tended to decrease while that of *pax7b* remained stable until day 30 post-injury (Figs 4A, 4b, 4C).

**Fig 4.**
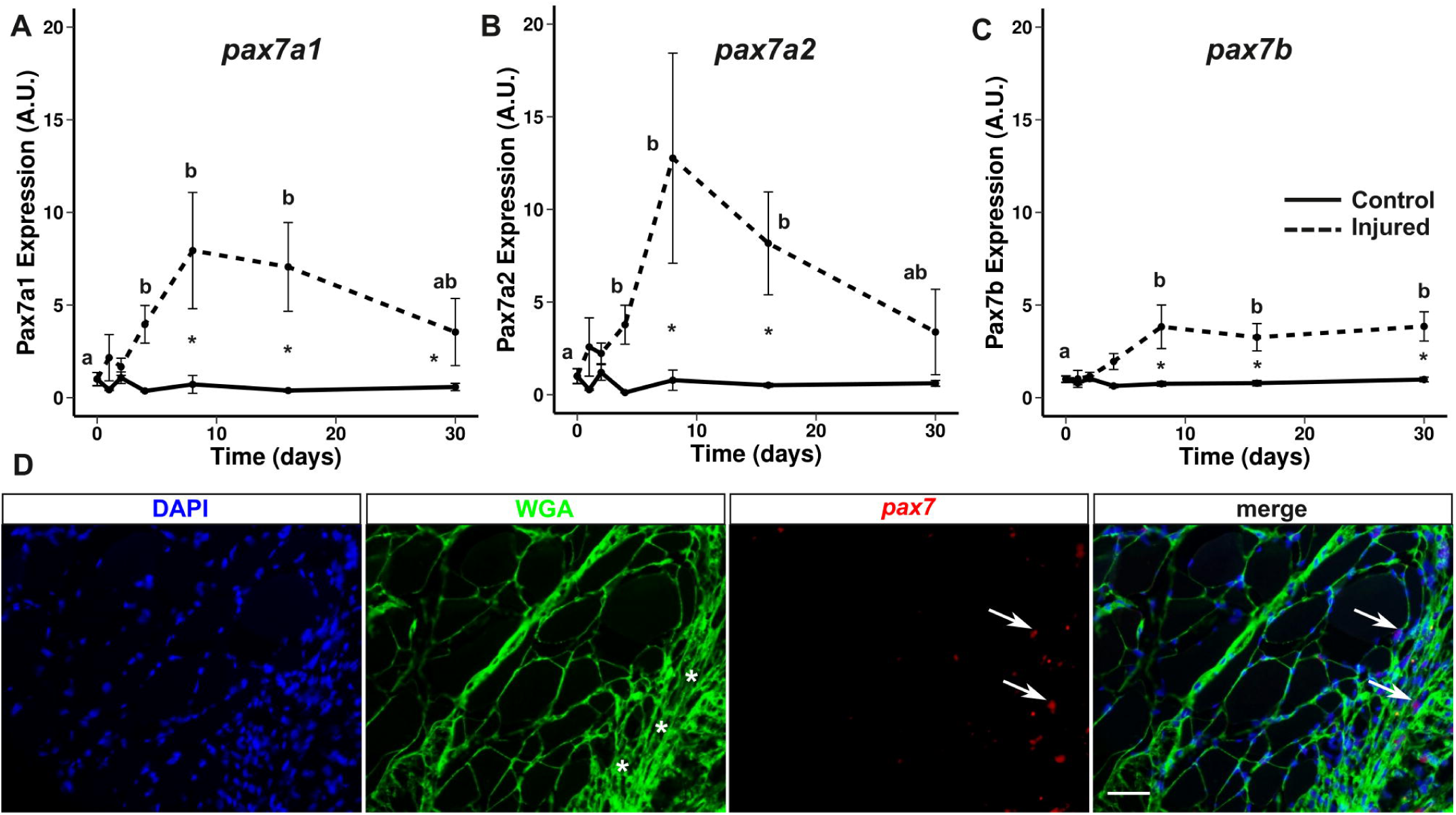
Expression of *pax7* genes increases during white muscle regeneration in trout. Gene expression profile of *pax7a1* (A), *pax7a2* (B) and *pax7b* (C) during muscle regeneration in rainbow trout normalized with *eF1a* expression. Bars represent the standard error and the letters indicate the significant differences between means within the same treatment (control or injured). The asterisk indicates significant differences between treatments at a given point. Statistical significance (p < 0.05) was determined by the Kruskal-Wallis rank test followed by the Dunn test. Localization of *pax7*^+^ cells (red) in 18-day injured muscle was performed by *in situ* hybridization (D). The extracellular matrix was stained with wheat germ agglutinin (WGA, green) and the nucleus was stained with DAPI (blue). The asterisks indicate the injury site and the scale bar corresponds to 50 µm.

To determine whether *pax7*^+^ cells accumulate at the site of injury during regeneration, we performed *in situ* hybridization for *pax7* on sections of regenerated muscle at day 18 (Fig 4D). At the site of injury, we observed accumulation of nucleus (DAPI) and extracellular matrix (WGA labeling). Consistent with qPCR results, *pax7*^+^ cells accumulated at the site of injury in contrast to the uninjured muscle area, where few *pax7*^+^ cells were observed.

### The proportion of Pax7^+^ cells decreases during in vitro differentiation of myogenic cells

To determine the evolution of the proportion of *pax7*^+^ cells during *in vitro* differentiation, we performed *in situ* hybridization for *pax7* and *myomixer* (*mmx*) on cultured myogenic cells. Fig 5A shows the presence of mononucleated cells expressing *pax7* and few cells expressing *mmx* at day 2 of culture. At day 7, we observed the presence of multinucleated myotubes with a strong expression of *mmx* and mononucleated cells expressing *pax7*^+^. Surprisingly, at day 7 we observed few cells expressing both *pax7* and *mmx,* although the intensity and the number of red dots (*pax7* labeling) were lower compared to what was observed at day 2. Quantification of the proportion of *pax7*^+^, *mmx*^+^ and *pax7*^+^/*mmx*^+^ cells is shown in Fig 5B. On day 2, the culture was composed of 44% *pax7*^+^ cells, 5% of *mmx*^+^ cells and 6% of *pax7*^+^/*mmx*^+^ cells. The percentage of *pax7*^+^ cells decreased to 18% on day 7, while the percentage of *mmx*^+^ and *pax7*^+^/*mmx*^+^ cells increased up to 19% and 31%, respectively.

**Fig 5.**
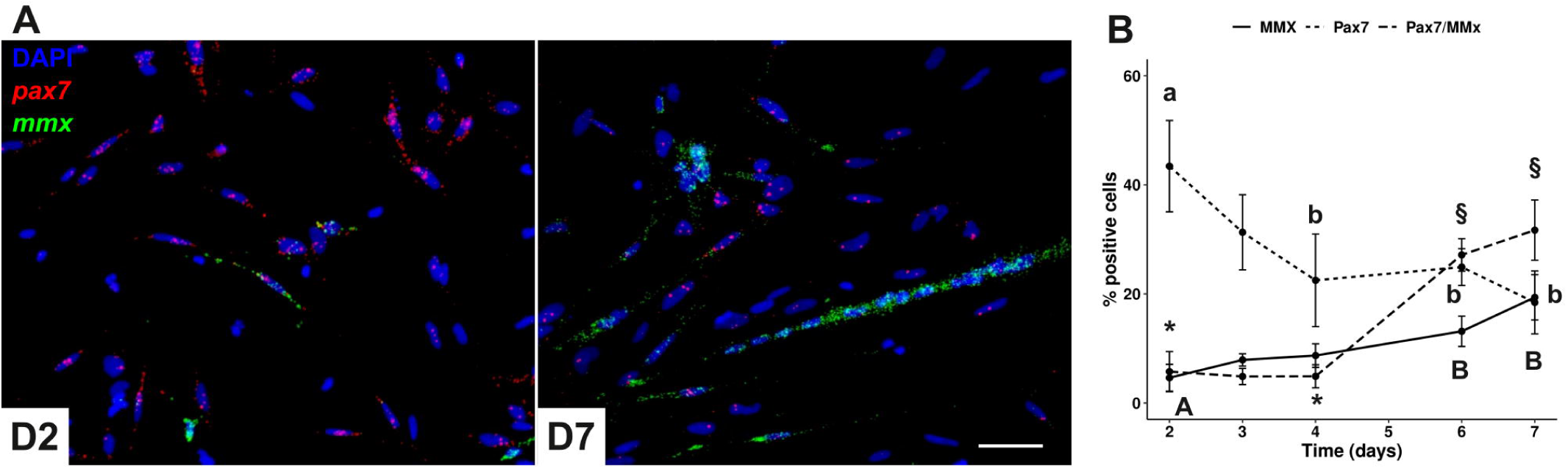
*Pax7* positive cells decrease during *in vitro* differentiation of myogenic progenitors. Myogenic progenitors were cultured in culture medium (DMEM, 10% SVF) for 7 days. Illustrative images of *pax7* (red) and *mmx* (green) *in situ* hybridization results performed at day 2 and day 7 of the culture (A). The scale bar corresponds to 50 µm. The percentage of *pax7* and *myomixer (mmx)* positive cells was determined by double *in situ* hybridization (B). Different capital letters indicate significant differences between means values of *mmx* positive cells. Different lowercase letters indicate significant differences between means of *pax7* positive cells. Different symbols indicate significant differences between means of *pax7/mmx* positive cells. Statistical significance (p < 0.05) was determined by Kruskal–Wallis rank test followed by Dunn’s test. The scale bar corresponds to 50 µm.

## Discussion

Adult muscle stem cells (satellite cells, *pax7*^+^ cells) are essential for fiber growth and muscle regeneration. Although the existence of these cells has been demonstrated in fish, the lack of highly sensitive and specific tools to label satellite cells has limited the study of their function. The aim of this work was to describe the dynamics of satellite cells (*pax7*^+^) during embryonic and larval stages, regeneration and *in vitro* differentiation in rainbow trout.

In most fish, the observation of satellite cells in adult muscle is difficult due to their small size and number, and the available tools are not sensitive and specific enough. In this context, we decided to take advantage of the recent improvement in *in situ* hybridization sensitivity using the novel technology RNAScope^®,17^, to localize *pax7*^+^ cells in trout muscle. In this species, we have previously identified three pax7 genes (*pax7a1*, *pax7a2* and *pax7b*) resulting from the salmonid-specific whole-genome duplication^20,24^. The coding sequences of these 3 genes are highly similar, sharing more than 88% of sequence identity. Given this high sequence identity, our probe set is able to detect the three *pax7* mRNAs as shown in Supplementary Fig 1, and thus all the cells expressing at least one *pax7* gene. Using this method, we were able to observe a *pax7* signal in embryos, white muscle and isolated satellite cells with no background (see Supplementary Fig 2).

In fish, *pax7* expression has previously been detected by classical *in situ* hybridization in the dermomyotome of zebrafish^25^ and trout^16,26^ until to the end of somitogenesis. Using RNAScope^®^ technology, our results confirmed the presence of *pax7*^+^ cells at the periphery of the myotome corresponding to the dermomyotome at day 14, assessing the specificity of the *pax7* probes. At the end of segmentation (D19-D21), *pax7*^+^ cells gradually appeared in the deep myotome from day 24 post fertilization. In addition, *pax7*^+^ cells were observed scattered throughout the somite, suggesting that *pax7*^+^ cells directly migrated directly from the dermomyotome to the deep myotome by moving between superficial muscle fibers. Thus, our results support the data obtained in zebrafish^27^ and brown trout^28^ showing that *pax7*^+^ cells also migrate between muscle fibers and not around their ends. Double labeling of *pax7* and the basal lamina, indicated that in trout, *pax7*^+^ cells are located beneath the basal lamina from day 112 post-fertilization, in line with observations in zebrafish, where *pax7*^+^ cells are also surrounded by the basal lamina as early as 6 days post-fertilization ^7,12^. In addition, *in situ* hybridization for *pax7* performed in muscle sections clearly showed the presence of *pax7*^+^ cells scattered in white muscle in juveniles (D136, ∼9g), in agreement with with data obtained with heterologous antibody in salmon^29^ but in contrast to zebrafish, where very few *pax7*^+^ cells are detected in white muscle^7^. This difference may be due to the persistence of a high rate of muscle hyperplasia in salmon in contrast to zebrafish^30^, which should require a high number of *pax7*^+^ cells.

In mammals, satellite cells are known to be required for muscle hyperplasia and hypertrophy, but also for muscle regeneration after injury. In trout, we have previously shown that mechanical injury induces the resumption of myogenesis as evidenced by the upregulation of *myogenin*, *myomaker* and *myomixer* and the formation of new small fibers^31,32^. The present results showed that during muscle regeneration the three *pax7* genes (*pax7a1, pax7a2* and *pax7b*) showed a peak of expression 8 days after injury, well before the peak of expression of *myogenin* and the formation of new fibres (D30)^31^. In addition, *in situ* hybridization with *pax7* probes at day 18, showed a large number of *pax7*^+^ cells at the wound site compared to the uninjured area. These results are consistent with previous work in adult^7^ and larval stage^12^ zebrafish showing that white muscle regeneration requires activation and proliferation of *pax7*^+^ cells. These results strongly suggest that in trout *pax7*^+^ cells are required for muscle regeneration and that the three *pax7* genes are involved. Indeed, in zebrafish, the two *pax7* genes (*pax7a* and *pax7b*) have been shown to have distinct functions, whereas in trout the specific roles, if any, of the three *pax7* genes remain unknown.

In mammals, *pax7* has been shown to be mainly expressed in quiescent and activated satellite cells^2^. Our results showed that 2 days after muscle cell extraction, 50% of the cells expressed *pax7* (44% *pax7*^+^ and 6% *pax7*^+^/*mmx*^+^) and were thus myogenic progenitors. In agreement with this result, using an antibody against MyoD, we observed that 60-70% of the extracted muscle cells were positive for MyoD^18^. The small difference may be due to the different markers used and to the fact that the trout in the present study are heavier (250g *versus* 5-10g). During the cell culture, the percentage of *pax7*^+^cells decreased from 44% to 18% on day 7, while the percentage of *mmx^+^* cells increased from 6% to 19%, in agreement with previous qPCR data^32^. Thus, as expected, the differentiating cells down regulated the expression of *pax7* and up regulated *mmx,* a differentiation marker. Surprisingly, the percentage of cells positive for both *pax7* and *mmx* increased from 5% on day 2 to 30% on day 7 of the culture. Close examination of the images, revealed that the *pax7* signal was weaker (fewer dots and lower intensity) in the *mmx*^+^ cells including myotubes. This result is reminiscent of previous observations in zebrafish showing the presence of myogenic cells expressing both *pax7* and *myogenin*^33^. Taken together, these results show that *pax7* expression occurs mainly in undifferentiated cells, although weak pax7 expression is still detectable in some differentiated cells.

In conclusion, the *in situ* RNAScope^®^ technology allowed us to localize the *pax7*^+^ cells with high sensitivity and specificity. We showed that *pax7*^+^ cells migrate into the deep myotome at the end of segmentation and reach their niche after hatching. In addition, we observed an accumulation of *pax7*^+^ cells at the wound site, suggesting their requirement for muscle regeneration and that *pax7* expression decreased in differentiating myogenic cells. Further work is needed to understand the effect of experimental conditions (fasting, aging, temperature) on the number of satellite cells.

## Acknowledgments

We particularly thank A Patinote and C. Duret for trout rearing and egg production and L Goardon of the fish facility PEIMA (Pisciculture Expérimentale INRAE des Monts d’Arrée) for muscle regeneration experiments.

---

**Supplemental Fig 1. The pax7 probe set enables simultaneous detection of all three *pax7* genes.** We developed a dot-blot hybridization approach in order to determine if the *pax7* probe set is able to target the three *pax7* genes present in the trout genome (*pax7a1*, *pax7a2*, *pax7b*) (Bio-Techne, Om-pax7b-cust # 575461). First, we transcribed cRNA sens of *pax7a1*, *pax7a2* and *pax7b* from *pax7* synthesized genes (Eurofins Genomics), using appropriate polymerase (Proméga, Riboprobe Systems sp6, # P1420, Riboprobe Systems T7, # P1440). The length of the cRNA is about 900 nt. We then applied 1µl (50ng) of each *pax7* cRNA onto a nitrocellulose membrane (Macherey-Nagel) and performed a dot-blot hybridization analysis. After UV fixation (UV crosslinker, Appligene Oncor), blots were incubated for 1 hour at 40°C with 1 drop of *pax7* probe set in order to hybridize with cRNA targets. After washing away excess probe, the presence of each *pax7* cRNA was detected using a chromogenic RNAscope kit (Bio-Techne, RNAscope 2.5 HD Reagent Kit – RED, # 322350). Red dots indicate that the *pax7* probe set recognizes the three *pax7* and doesn’t recognize the negative control corresponding to the *pax7a1* antisense RNA.

**Supplementary Fig 2. Specific pax7 signal observed in white muscle section and in vitro myogenic cells** Transverse sections (A, B) of trout white muscle (10g) were analyzed by *in situ* hybridization for *pax7* (red) and extracellular matrix was stained with Alexa 488-conjugated wheat germ agglutinin (green). Arrowheads indicate *pax7*^+^ cells beneath the basal lamina of the muscle fibers. No signal was observed with the negative probe against a bacterial gene *DapB* (B).

Myogenic progenitors (C, D) cultured for 1 day in culture medium (DMEM, 10% SVF) were analysed by *in situ* hybridization for *pax7* (red). A strong signal was observed in mononucleated cells (C). No signal was observed with the negative probe against a bacterial gene *DapB* (D). The nuclei are counterstained with DAPI and the scale bar corresponds to 50 µm.

