## Supplementary figures and images for "Dynamics of pax7 expression during development, muscle regeneration, and *in vitro* differentiation of satellite cells in the trout"

### Supplemental Figure 1

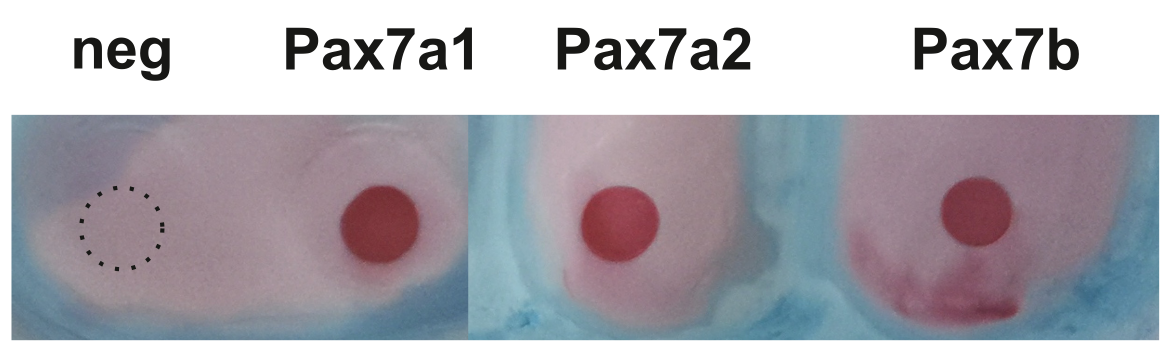

### Supplemental Figure 2

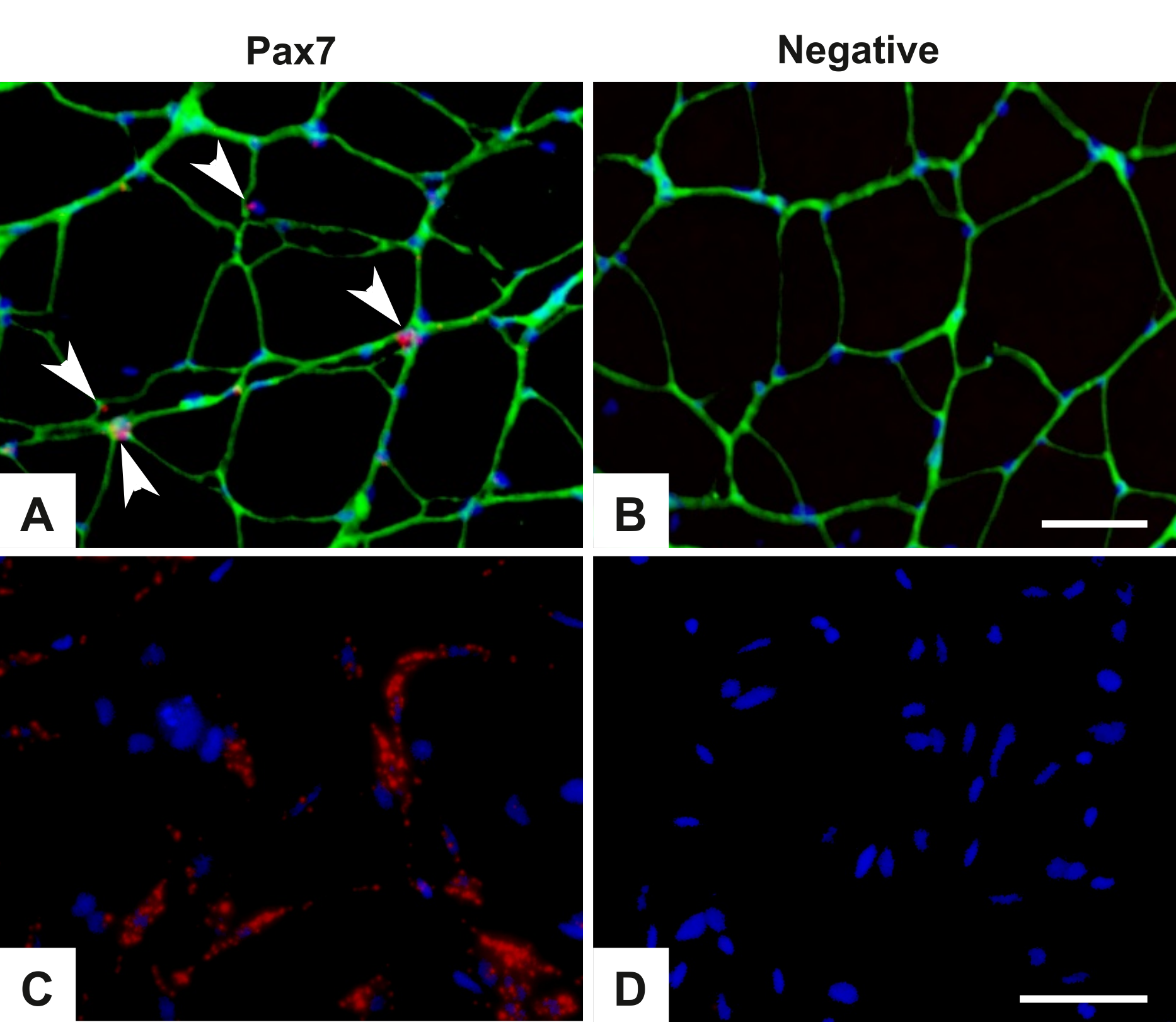
